# Perch selection of three species of kingfishers at the Pantanal wetland, Brazil

**DOI:** 10.1101/2020.09.21.306027

**Authors:** Laura C. Peinado, Zaida Ortega

## Abstract

Animal movement and behavior depend on the distribution of resources on the habitat. Therefore, individual animals are constantly making decisions on resource selection based on different attributes of the resource or its associated environmental variables. For fish-eating birds as kingfishers, selecting a suitable perch can report many benefits, as improving fishing success or reducing predation risk. Nowadays, not only natural structures, as branches, are available for birds to perch but also artificial ones, as electric lines. Thus, we aimed to understand which variables drive kingfishers’ perch selection, including the potential effect of its anthropic origin. We studied perch selection of three species of kingfishers inhabiting the Pantanal of Miranda of Brazil: *Megaceryle torquata, Chloroceryle amazona* and *Chloroceryle americana*. They feed in temporary ponds that are rich in trophic resources, where they have both natural and artificial potential perches. We hypothesized that artificial perches could be strongly selected, as they are more stable than natural ones and go through the ponds, providing a long surface to select optimal conditions. We assessed how kingfishers are selecting perches based on four ecologically relevant traits: (1) being artificial or natural, perch height, (3) distance to the water, and (4) plant cover. We used a resource selection function (RSF) approach to quantify the effect of these variables in the probability of presence of kingfishers. The artificial origin of a perch was independent of the probability of selection for the three species. Furthermore, birds acted randomly to the other studied variables, except for individuals of *C. amazona*, which select higher perches, above 3.20 m. We discuss the implications of these results for understanding the behavioral ecology and use of space of neotropical kingfishers, and how this affects their vulnerability to human habitat alterations.

**GRAPHICAL ABSTRACT:** 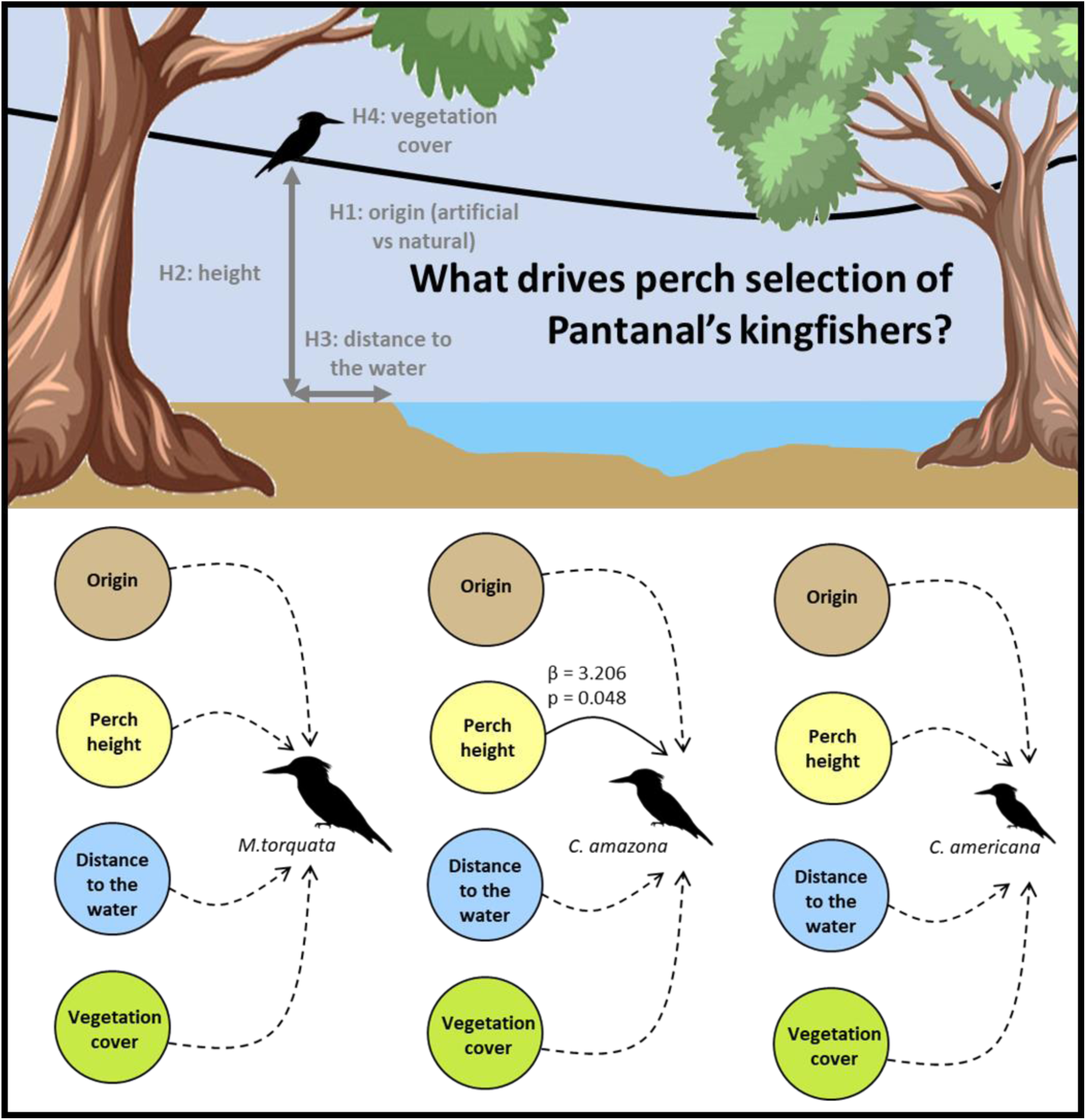

## INTRODUCTION

The habitat includes the set of conditions and resources necessary to support a population in space and time (McComb 2016). These conditions and resources are required for organisms to live and, at the same time, drive the spatial distributions of individuals within the habitat (Manly et al. 2007). Consequently, the spatial distribution of animals is not random, but it is modulated by how resource availability influences survival and reproduction (Litvaitis and Villafuerte 1996). For birds, the structure and composition of the vegetation and water bodies are key factors, since they drive the diversity and abundance of trophic resources, perches, and nesting sites (Cueto 2006).

The amount and quality of resources vary in space and time, conditioning the behavioural decisions of animals. Animals tend to select the microhabitats that maximize their fitness (Chalfoun and Schmidt 2012), depending on their specific needs and balancing the associated -energetic and non-energetic-costs and benefits (Hutto 1985; Block and Brennan 1993; Houston and Lang 1998; Johnstone and Earn 1999). Hence the importance of habitat selection as the main driver of animal choices (Cody 1985). It is expected that animals select –that is, use above the available– the microhabitats where they achieve higher reproductive success and survival (Levins 1968, Orians 1980). At the same time, animals should avoid –that is, use below the available– those places where chances of reproduction and survival are low (Orians and Wittenberger 1991). Here lays the importance of studying how and why animals select the different available resources (Cody 1985). Moreover, the answer to the question ‘why animals select a certain habitat?’ is important to learn about behavioral and evolutionary ecology and apply it to manage and preserve their relevant habitats (Chalfoun and Schmidt 2012).

Certain structures, as perches, are known to be important habitat elements for birds, since they are related to multiple ecological functions (Cody 1985). For example, birds use perches for defending their territory (Wiens 1969), reproduction (Krams 2001), predating (Wolff et al. 1999), breeding and feeding (Glinski and Ohmart 1983), and even resting. Due to their relevance for many vital processes of birds, selecting suitable perches may entail important ecological consequences. For example, perch selection of some species drives seed dispersion and its consequent ecological succession (Athiê and Dias 2016, Vogel et al. 2016). For fish-eating birds, microhabitat use is closely related to the temporal and spatial distribution of water bodies (Ferreira 2013). There is evidence that these birds use microhabitats with higher fish abundance (Elmberg et al. 2010) and, accordingly, one would expect them to select perches that enhance fishing success. An important group of continental fish-eating birds are kingfishers. Three species of kingfishers: *Megaceryle torquata* (Linnaeus 1766), *Chloroceryle amazona* (Latham 1790) and *Chloroceryle americana* (Gmelin 1788) are particularly abundant at the Pantanal wetland. They are syntopic and differ in body size, *M. torquata* (≈ 39.6 cm of mean body length) > *C. amazona* (≈ 29.5 cm) > *C. americana* (≈ 21.8 cm) (Gwynne et al. 2010, Rodrigues et al. 2019). The three species inhabit temporal ponds of the Pantanal, where they share the habitat and diet. They mainly feed on shallow water fish although they can occasionally eat arthropods, aquatic insects, crustaceans and even lizards. At these temporal ponds, perches serve to visualize prey and prepare the attack from a certain height (Gwynne et al. 2010).

The Pantanal is a natural reserve, holding one of the worldwide highest abundance of birds (Tubelis and Tomas 2003). In the Anthropocene, however, artificial elements, as electric lines, are present even at the most isolated natural reserves. These artificial structures are then amongst the available options for fish-eating birds to use as perches, where, in fact, they are commonly observed (pers. obs.). Electric lines may offer a better structure to visualize prey, free of the visual obstacles that branches usually present, particularly in tropical areas. However, a bird perching in a line would also be more exposed to potential predators, and even experience risk of electrocution. Our study aims to understand perch selection of kingfishers and to elucidate the role of electric lines on it. We hypothesized that electric lines could be selected over natural perches (Figure 1; H1), since they could offer benefits for kingfishers to enhance fishing success. In addition, we hypothesized that other attributes of the perches, such as height (H2), distance from the water (H3) and vegetation cover (H4) may influence perch selection of kingfishers at the Pantanal (Figure 1). Differences in perch selection could arise from differences in body size and, maybe, behavior of the different species, and would also contribute to reduce interspecific competition. However, we did not have a priori expectations about the direction of the differences in perch selection of the three species.

**Figure 1.**
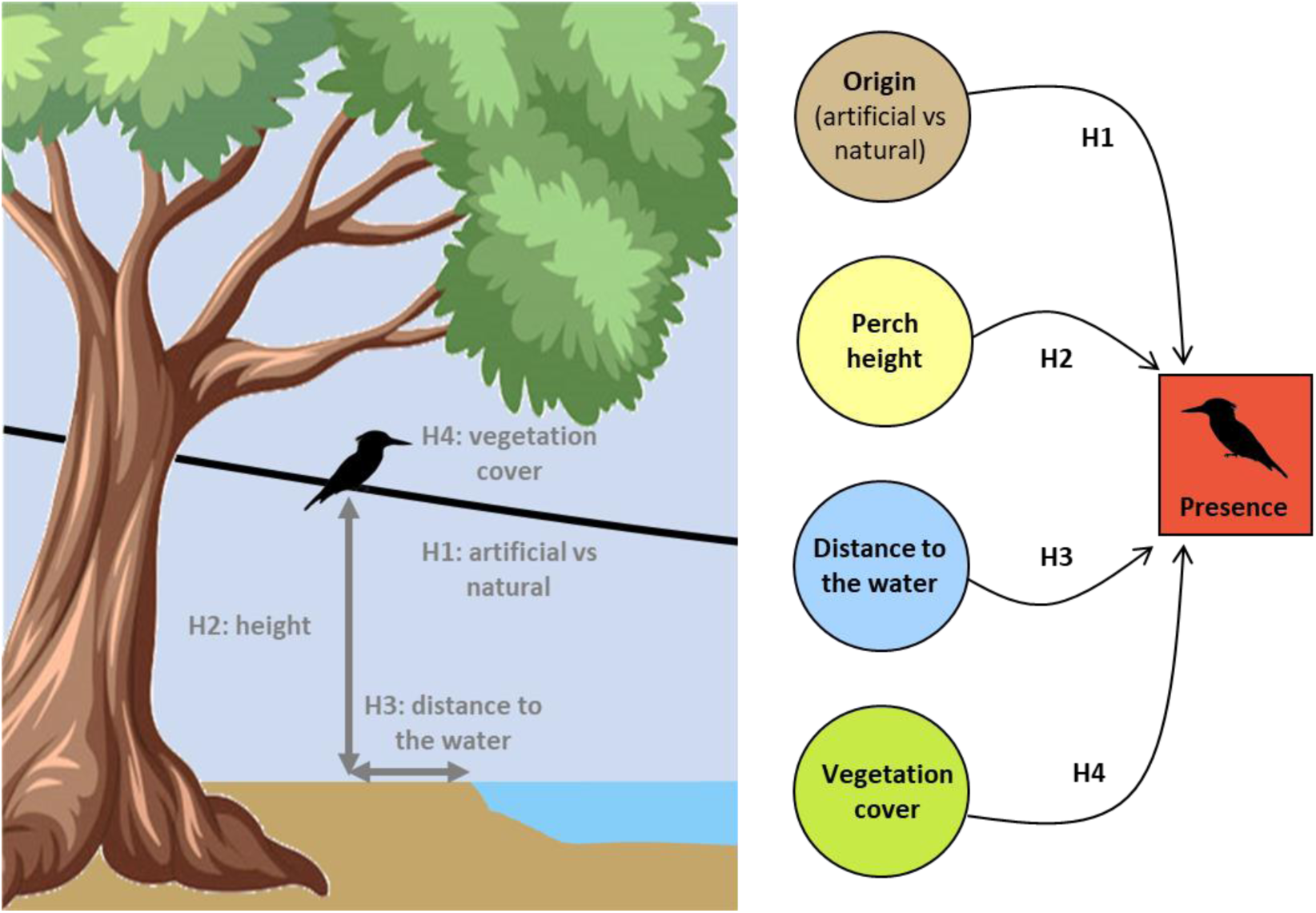
Hypotheses on perch selection of the three species of kingfishers (*Megaceryle torquata, Chloroceryle amazona*, and *C. americana*) studied at the Pantanal of Brazil: (H1) birds of the three species will select artificial over natural perches; perch height (H2), distance to the water (H3) and vegetation cover above the animal (H4) will affect perch selection, probably in a different way for the different species.

## METHODS

### Study area

Our study took place at the Pantanal of Miranda (Corumbá, Mato Grosso do Sul, Brazil) (19°34’ S, 57°01’ W). The Pantanal is one of the biggest wetlands of the world, with approximately 150.000 km^2^, and is characterized by flooded grasslands and savannas. Climate is tropical, with marked dry and wet seasons and a mean annual temperature of 25 °C (Bergier and Assine 2016). We conducted fieldwork in September of 2018, during the dry season. We sampled 13 temporal ponds located along the MS-184 track that crosses the Pantanal Matogrossense National Park at the municipality of Corumbá. These temporal ponds are quite similar: they are well preserved, maintaining the native vegetation and highly rich and abundant in fish (Alho 2008). Despite the place is well preserved, each pond has some artificial elements, typically: a crossing wood bridge, a simple electric line and informative signals, which are the unique artificial elements there.

As we did not capture or marked the individuals, we sampled the ponds sequentially, sampling each pond just once, in order to avoid pseudoreplication. All observations were conducted in the period of maximum activity of kingfishers, between 7.00 and 11.00 h. We visited the ponds and registered four variables of the perch where an individual kingfisher was posed (used perch) and the closest four available (non-used) perches. We defined the available perches as the closest projection, whether it was artificial or natural, to the perch used by the bird. By registering those variables simultaneously for each used/available perch for each individual bird, we prevented temporal and environmental variation that would confound results. The studied variables were: (1) artificial vs natural perch, (2) perch height, distance from the water, and (4) vegetation cover. Due to the difficulty of approaching the perches, we estimated their height and distance to the water by eye, always by the same previously trained observer (LCP). Height was estimated as the vertical distance (in m) between the perch and the floor (whether is land, grass, or water). Distance to the water was estimated as the horizontal distance (in m) to the pond shore, considered zero if the perch was exactly at the shore, negative if it was within the pond border and positive if it was outside the pond border. We quantified vegetation cover as the percentage of vegetation at the angle of view of the observed kingfisher, dividing the plane into four quadrants to facilitate the estimation (Figure 1). We observed a total of 31 birds using perches at these ponds: 15 *M. torquata*, 14 *C. amazona* and 5 *C. americana*, being each used perch associated with their respective 4 non-used (available) perches (60 *M. torquata*, 56 *C. amazona* and 20 *C. americana* non-used perches).

### Statistical analyses

In order to analyze how kingfishers select their perches, we used a resource selection function (RSF) approach. A RSF is a function of the probability of an individual using a specific resource based on its availability in the habitat (Manly et al. 2007). In our case, the resource is the perch and we are interested in the effect of the four studied variables in their probability of use. Here we used mixed conditional logistic regression (CLR) to solve the RSF, pairing the data of used and available perches for each individual bird (stratum). This way we know that available perches are true absences that each individual could had selected in the moment of observation but did not do so (Jones 2001; Duchesne et al. 2010; BenÍcio et al. 2019). The response variable is the presence of the bird in the perch (1 = used perch / 0 = unused perch). This way, with CLR we model the probability of a bird selecting a perch in relation with other available perches depending on the effect of the explicative variables (artificial/natural perch, height, distance to water and vegetation cover) in the same probabilistic process (Liedke et al. 2018, Ortega et al. 2019). We used the function clogit of the package ‘survival’ (Therneau 2015) of R (R Core Team 2018). As the CLR model did not converge when the species were considered together (that is, including the interaction between the variable ‘species’ and the four fixed factors), we fitted a CLR model to each studied species.

## RESULTS

The three species used perches with quite similar traits, at approximately 1-4 m height, at 2-3 m within the pond shore (in horizontal, see Figure 1), with 5-20 % of vegetation within the bird’s viewpoint (Table 1). In general, we obtained that these three kingfisher species used artificial perches in a proportion of 8/25 artificial/natural perches, a similar proportion as available over the pond. In addition, results from the CLR revealed that *M. torquata* used the perches independently of being artificial or natural, their height, horizontal distance to water shore or vegetation cover (Table 2). Individuals of *C. amazona* selected perches higher than the mean availability, specifically they selected perches higher than 3.20 m (Figure 2), but independently of the distance to water, being artificial or natural or the vegetation cover (Table 2). Within the range of perches studied (∼ 0 to10 m height), the probability of finding a *C. amazona* individual increases in almost 2.5 times for each meter of altitude. For *C. americana* sample size was small, so we decided to only include one variable in the model. Since distance to the water shows the biggest mean difference between used and available places for this species (Table 1), we fitted the effect of the distance to the water in probability of presence. However, individuals of *C. Americana* selected their perches independently of its distance to the water shore (Table 2).

**Table 1.** Mean and standard deviation values of the studied variables for used and available perches of the three kingfisher species studied at the Pantanal of Miranda (Mato Grosso do Sul, Brazil).

| Variable | Perch | *M. torquata* |  | *C. amazona* |  | *C. americana* |  |
| --- | --- | --- | --- | --- | --- | --- | --- |
|  |  | Mean | SD | Mean | SD | Mean | SD |
| Perch height (m) | Used | 4,08 | 2,55 | 2,91 | 2,53 | 1,38 | 0,22 |
|  | Available | 3,23 | 2,55 | 1,73 | 1,34 | 1,64 | 1,01 |
|  | Total | 3,39 | 2,55 | 1,96 | 1,69 | 1,59 | 0,90 |
| Distance to water (m) | Used | -2,12 | 3,14 | -2,73 | 3,00 | -3,05 | 2,50 |
|  | Available | -2,32 | 3,83 | -2,32 | 3,39 | -2,44 | 2,22 |
|  | Total | -2,28 | 3,69 | -2,40 | 3,30 | -2,56 | 2,23 |
| Vegetation cover (%) | Used | 6,54 | 17,00 | 9,29 | 14,78 | 0,00 | 0 |
|  | Available | 8,37 | 17,02 | 9,38 | 16,21 | 4,69 | 5,61 |
|  | Total | 8 | 16,90 | 9,35 | 15,83 | 3,75 | 5,34 |
| Artificial | Used | 5/13 |  | 3/14 |  | 0/4 |  |
|  | Available | 34/52 |  | 21/56 |  | 0/16 |  |
|  | Total | 23/65 |  | 24/70 |  | 0/20 |  |

**Table 2.** Results of the mixed conditional logistic regression models for the probability of use of perches by the three species of kingfishers (*Megaceryle torquata, Chloroceryle amazona*, and *C. americana*), studied at the Pantanal of Brazil, based in the four studied variables (perch height, distance from water, vegetation cover and artificial/natural perch).

| Species | Variable | Coefficient | Exp (coefficient) | SE | Z | p-value |
| --- | --- | --- | --- | --- | --- | --- |
| *M. torquata* | Perch height (m) | 0.411 | 1.508 | 0.263 | 1.560 | 0.119 |
|  | Distance to water (m) | 0.194 | 1.214 | 0.214 | 0.908 | 0.364 |
|  | Vegetation cover (%) | -0.019 | 0.981 | 0.029 | -0.631 | 0.528 |
|  | Artificial | 0.603 | 1.828 | 1.353 | 0.446 | 0.656 |
| *C. amazona* | Perch height (m) | 3.206 | 2.469 | 1.625 | 1.974 | **0.048** |
|  | Distance to water (m) | -1.710 | 8.428 | 2.996 | -0.571 | 0.568 |
|  | Vegetation cover (%) | 8.186 | 1.085 | 6.869 | 1.192 | 0.233 |
|  | Artificial | -1.912 | 4.985 | 1.895 | -0.001 | 0.999 |
| *C. americana* | Distance to water (m) | -1.275 | 0.279 | 1.407 | -0.907 | 0.365 |

Model *M. torquata*: Likelihood ratio test = 4.17, 4 df, p = 0.4
Model *C. amazona*: Likelihood ratio test = 21.08, 4 df, p < 0.001
Model *C. americana*: Likelihood ratio test = 1.66, 1 df, p = 0.2

**Figure 2.**
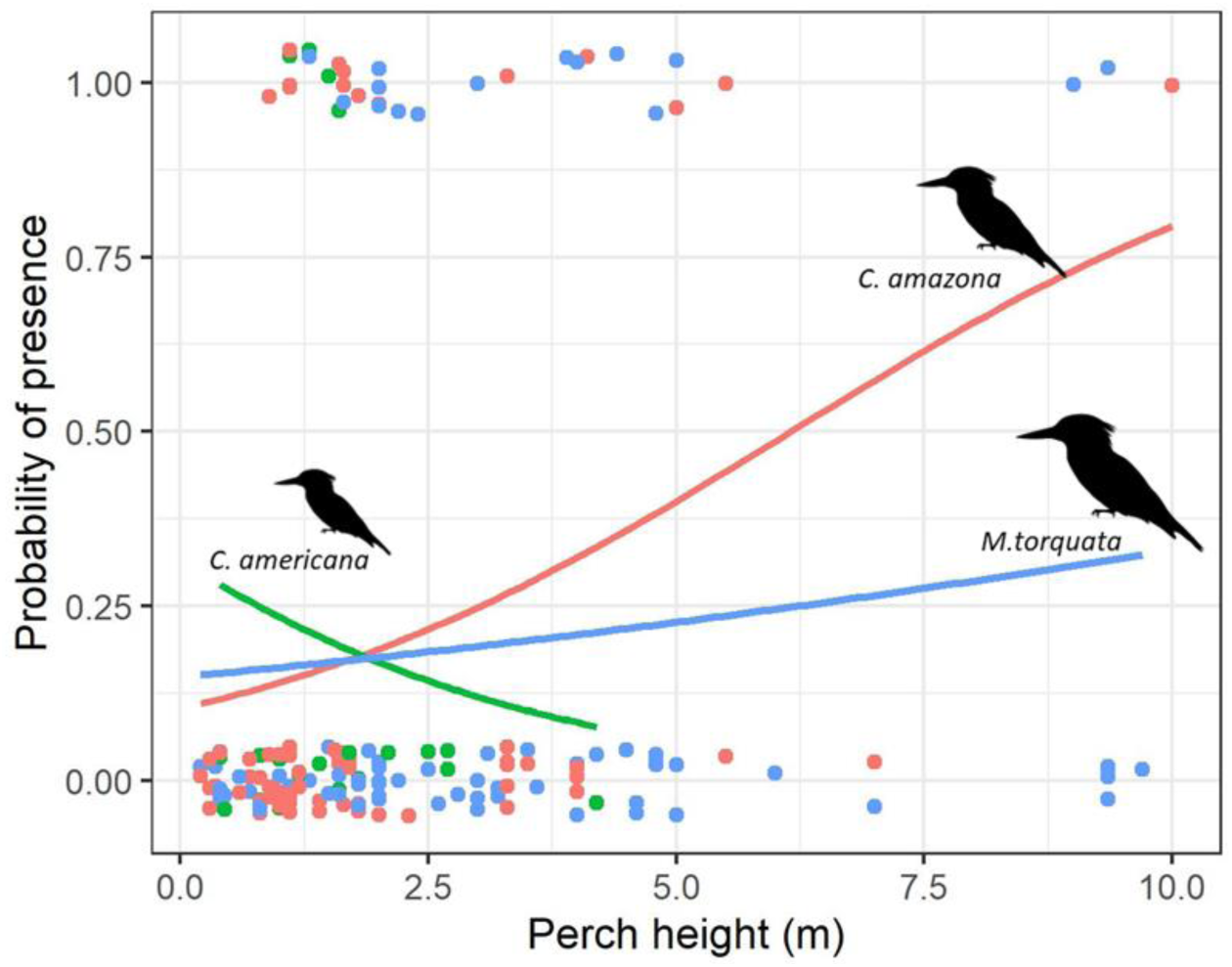
Probability of presence depending on perch height for the three studied species, estimated by a logistic regression model. Influence of perch height on the presence of birds was only significant for *Chlorocetyle amazona* (see more details in the text).

## DISCUSSION

This is the first study on habitat selection of kingfishers using a resource selection function approach. We assessed perch selection of three syntopic species of kingfishers living at temporal ponds of the Pantanal wetland, aiming to understand the role of relevant habitat covariates and, particularly, whether birds were selecting artificial perches (such as electric lines) over natural ones. Our approach -assessing the traits of used perches in comparison with the available perches for each bird within a close temporal and spatial scale-allowed us getting robust conclusions on habitat selection, precluding confounding effects. One species (*C. amazona*) select perches higher than the mean available in the habitat while acting randomly regarding distance to water, vegetation cover and the fact of being natural or artificial. The other species (*M. torquata* and *C. americana*) do not select their perches based on any of the studied traits.

Contrary to our expectation, kingfishers from the Pantanal are randomly using artificial and natural perches. The three studied species, *M. torquata, C. amazona* and *C. americana*, use both artificial perches (bridges and electric lines) and natural perches (tree branches) in the same relative proportion as available in their habitat. This result suggests that artificial structures ─at least at this isolated natural reserve─ are not changing the behaviour of these fish-eating birds. As kingfishers do, raptors also use perches for hunting, feeding and resting (Reinert 1984). Perching minimizes the required energy budget for hunting, feeding and resting when compared with flying and hovering (Collopy and Koplin 1983). The studied kingfishers used artificial and natural perches randomly, which suggests that their energetic balance of costs and benefits would be similar. Moreover, artificial perches add to the natural perches available for kingfishers, increasing the available places to hunt or rest, and could even lead birds to reach some places that would be inaccessible otherwise (Hall et al. 1981).

Other birds are known to use artificial perches to save energy in their feeding behavior (Sheffield et al. 2001). However, there are also cases when the introduction of artificial structures modify the behavior of birds, as with the introduction of artificial lights (Longcore 2010, Kempenaers et al. 2010), or noisy elements (Polak 2014). We think that the fact that the studied kingfishers use natural or artificial perches randomly may reflect that both perch types offer similar access to food, minimizing the energy necessary for fishing (Resende 2008). Furthermore, as the region is quite isolated and well-preserved, we think that the little artificial structures there entail a low level of alteration or disturbance in the ecology of kingfishers (Junk et al. 2006).

The three kingfisher species use similar perches, situated at ∼ 1-4 m height and 2-3 m of horizontal distance within the pond border. They pose in perches with a vegetation cover of 5-20 % at their viewpoint. Approximately, the 75 % of the perches used by Pantanal’s kingfishers are natural structures, as tree branches, being the other 25 % artificial structures, as energy lines and traffic signals. In the literature, we found that *M. torquata* perches at variable or high perch height, *C. amazona* at low height and exposed perches and *C. americana* always at low perches (Gwynne et al. 2010). Our results showed that only *C. amazona* select for height, perching higher than 3.20 m. For two species of kingfishers studied in Japan, differences in perching height, streaming velocity and prey sizes were found (Kasahara and Katoh 2008). This study confirmed the differentiation of niche in these species and a positive relationship between body size and height of used perches in several cases (Willard 1985, Bonnington et al. 2008, Kasahara and Katoh 2008). Our study does not support this relationship, because the medium size kingfisher selected perches higher than the biggest kingfisher. Individuals of *C. amazona* select perches higher than 3.20 m, while birds of the other two species act randomly regarding perch height. Among the three kingfishers, *C. amazona* presents a medium body size. Thus, their selection of higher perches would not be related to their body size. The Pantanal -because of its concentration and abundance of wildlife-brings animals a context of prey abundance (Swarts 2000), particularly on fishes (Resende 2008). Despite this, *C. amazona* and *M. torquata* have similar diets and choose fishes with similar sizes (Willard 1985). Thus, it is possible that perch segregation may function to relax competition between these two species.

At the study area, the thirteen isolated little studied ponds are similar in depth, stream, turbidity, width. Sampling was conducted in the dry season, when streams are shallow and low, forming rounded little ponds close to wooden bridges of the track. It is important to provide this framework because the Pantanal is completely different during the wet season. Because of the similarity of water-related variables in the dry season, perch selection depends on other aspects of perches, as vegetation cover, distance to water, height and natural or artificial origin. At the ponds of the Pantanal, both predation risk and food availability would be similarly high for the available perches during the dry season (Holbrook and Schmitt 1988). A perch without vegetation cover and close to the water may enhance fishing success, while increasing predation risk. On the contrary, using a perch hidden among leaves may reduce predation risk but lower fishing success. In an environment of high availability of fishes and high predation risk as the Pantanal, it is possible that a trade-off between these two forces would lead birds to use perches quite randomly.

We conclude that Pantanal kingfishers selected their perches randomly regarding artificial/natural origin, distance to water, height above ground and vegetation cover. The exception was *C. amazona*, which selected higher perches, probably to avoid interspecific competition with *M. torquata*. We propose that the high food available and the similarity in predation risk, also high at the Pantanal, may result in similar quality of the available perches for kingfishers. Future studies assessing perch selection at different seasons or at places in a gradient of food availability/predation pressure will throw light in this hypothesis. We think that the isolation and good preservation of the Pantanal, with few artificial structures, may prevent the existing artificial structures, as electric lines, to modify perch selection for the moment. However, bushfires are dramatically increasing in the area, fueled by climate change and predicted to get worse in future years if policies and land use remain the same. More than 14.000 km^2^ of native vegetation are already predicted to be lost in the area until 2050 (). We think that *C. amazona* can be particularly affected by the loss of native vegetation related to bushfires and we recommend conducting extensive research on habitat selection of Neotropical birds to estimate the vulnerability of species and prioritize preserving the species selected habitat elements.

## ACKNOWLEDGMENTS

This article is the result of a research of the 21^st^ Field Course Ecology of the Pantanal (EcoPan2018) of the Federal University of Mato Grosso do Sul. A postdoctoral fellowship PNPD/CAPES (#1694744) supported Z.O. and a master fellowship CNPQ (#130842/2017-6) supported L.C.P. Our study was purely observational, nevertheless, fieldwork was approved by the Ethics Committee of the Federal University of Mato Grosso do Sul. The author’s contributions were: LCP and ZO conceived and designed the study, LCP and ZO conducted the research, LCP and ZO analyzed the data, LCP wrote the paper and both authors revised and edited the manuscript. Thanks to all colleagues, teachers and friends of the Ecopan 2018 that help in this research, particularly Ana Belén Ávila.

## LITERATURE CITED

Alho, C.J.R. (2008). Biodiversity of the Pantanal: response to seasonal flooding regime and to environmental degradation. Brazilian Journal of Biology 68:957–966.

Athiê, S., & Dias, M. M. (2016). Use of perches and seed dispersal by birds in an abandoned pasture in the Porto Ferreira state park, southeastern Brazil. Brazilian Journal of Biology, 76:80–92.

Benício R., Z. Ortega, A. Mencía, and D. Cunha-Passos (2019). Microhabitat selection of Ameiva ameiva (Squamata: Teiidae) in the Brazilian Pantanal. Herpetozoa 31:211–218.

Bergier, I., and M. Assine (2016). Dynamics of the Pantanal Wetland in South America. In The Handbook of Environmental Chemistry (D. Barceló, A.G. Kostianoy, Editors). New York: Springer International Publishing 243.

Block, W. M., and L. A. Brennan (1993). The habitat concept in ornithology. Theory and Applications. In Current Ornithology (D. M. Power, Editor). Plenum Press, NY, USA.

Chalfoun, A. D., and K. A. Schmidt (2012). Adaptive breeding-habitat selection: is it for the birds?. The Auk 129:589–599.

Cody, M.L. (1985). Habitat selection in birds. Academic Press, Florida, OR, USA.

Cueto, V.R (2006). Escalas en ecología: su importancia para el estudio de la selección de hábitat en aves. El hornero 21:1–13.

Duchesne, T., D. Fortin, and N. Courbin (2010). Mixed conditional logistic regression for habitat selection studies. Journal of Animal Ecology 79:548–555.

Durán, A.A. (2017). Datos preliminares sobre la influencia de la turbidez del agua y profundidad en el éxito de captura de presas por Megaceryle torquata (aves, Alcedinidae). Revista Biodiversidad Neotropical 7:152–155.

Elmberg, J., L. Dessborn, and G. Englund (2010). Presence of fish affects lake use and breeding success in ducks. Hydrobiologia 641:215–223.

Ferreira R.P. (2013). Influência da altura do poleiro de ataque no sucesso de captura de Megaceryle torquata (Aves: Alcedinidae) no Pantanal do Miranda. In Ecologia do Pantanal – Curso de Campo (E.C. Corrêa, G. Gris, E. Sczeny-Moraes, et al., Editors). Editora UFMS, Campo Grande, Brazil.

Guerra, A., de Oliveira Roque, F., Garcia, L.C., Ochoa-Quintero, J.M., de Oliveira, P.T.S., Guariento, R.D., & Rosa, I.M. (2020). Drivers and projections of vegetation loss in the Pantanal and surrounding ecosystems. Land Use Policy 91:104388.

Gwynne, J.A., R.S. Ridgely, M. Argel, and G. Tudor (2010). Guia Aves do Brasil: Pantanal & Cerrado. Wildlife Conservation Society,Editora Horizonte, São Paulo.

Holbrook, S.J., and R.J. Schmitt (1988). The combined effects of predation risk and food reward on patch selection. Ecology 69:125–134.

Houston, A.I, and A. Lang (1998). The ideal free distribution with unequal competitors: the effects of modelling methods. Animal behaviour 56:243–251.

Hutto, R.L (1985). Habitat selection by nonbreeding migratory land. In Habitat selection in birds (M. Cody, Editor). Academic Press, London, UK.

Johnstone, R.A., and D.J. Earn (1999). Imperfect female choice and male mating skew on leks of different sizes. Behavioral Ecology and Sociobiology 45:277–281.

Jones, J. (2001). Habitat selection studies in avian ecology: a critical review. The auk 118:557–562.

Junk, W.J., C.N. Da Cunha, K.M. Wantzen, P. Petermann, C. Strüssmann, M.I. Marques, and J. Adis (2006). Biodiversity and its conservation in the Pantanal of Mato Grosso, Brazil. Aquatic Sciences 68:278–309.

Krams, I. (2001). Perch selection by singing chaffinches: a better view of surroundings and the risk of predation. Behavioral Ecology 12:295–300.

Levins, R. (1968). Evolution in changing environments: some theoretical explorations (No. 2). Princeton University Press. Princeton, New Jersey.

Liedke, A.M.R.., R.M. Bonaldo, B. Segal, C.E.L. Ferreira, L.T. Nunes, A.P. Burigo, S. Buck, L.G.R. Oliveira-Santos, and S.R. Floeter (2018). Resource partitioning by two syntopic sister species of butterflyfish (Chaetodontidae). Journal of the Marine Biological Association of the United Kingdom 98:1767–1773.

Litvaitis, J. A., and R. Villafuerte (1996). Factors affecting the persistence of New England cottontail metapopulations: the role of habitat management. Wildlife Society Bulletin 686–693.

Manly, B.F. L., L. McDonald, D.L. Thomas, T.L, McDonald, and W.P. Erickson (2007). Resource selection by animals: statistical design and analysis for field studies. Kluwer Academic Publishers, Dordrecht, Netherlands.

McComb, B.C. (2016). Wildlife habitat management: concepts and applications in forestry. CRC Press, Boca Raton, FL, USA.

Orians, G.H. (1980). Habitat selection: General theory and applications to human behavior. The evolution of human social behavior. Elsevier North Holland, New York.

Orians, G.H., and J.F. Wittenberger (1991). Spatial and temporal scales in habitat selection. The American Naturalist 137:S29–S49.

Ortega, Z., A. Mencía, K. Martins, P. Soares, V.L. Ferreira, and L.G.R. Oliveira-Santos (2019). Disentangling the role of heat sources on microhabitat selection of two Neotropical lizard species. Journal of Tropical Ecology 35:149–156.

R Core Team. (2018): R: a language and environment for statistical computing. R Foundation for Statistical Computing, Vienna, Austria.

Reinert, H.K. (1984). Habitat variation within sympatric snake populations. Ecology 65:1673–1682.

Resende, E.K.D.R. (2008). Pulso de inundação: processo ecológico essencial à vida no Pantanal. Embrapa Pantanal, Corumbá, MS, Brasil.

Rodrigues, R.C., É. Hasui, J.C. Assis, J.C.C. Pena, R.L. Muylaert, V.R. Tonetti, F. Martello, A.L. Regolin T.V. Viera da Costa, M. Pichorim, E. Carrano, et al. (2019). Atlantic bird traits: a dataset of bird morphological traits from the Atlantic forests of South America. Ecology 100:e02647.

Sheffield, L.M., J.R. Crait, W.D. Edge, and G. Wang (2001). Response of American kestrels and gray-tailed voles to vegetation height and supplemental perches. Canadian Journal of Zoology 79:380–385.

Swarts, F. A. (2000). The Pantanal in the 21st century: For the planet’s largest wetland, an uncertain future. In The Pantanal: understanding and preserving the world’s largest wetland (F.A. Swarts, Editor). Selected papers and addresses from the World Conference on Preservation and Sustainable Development in the Pantanal, Paragon House, St.Paul, Minn 1–22.

Therneau, T.M. (2015). A Package for Survival Analysis in S. version 2.38, https://CRAN.R-project.org/package=survival.

Tubelis, D.P., and W.M. Tomas (2003). Bird species of the Pantanal wetland. Brazil. Ararajuba 11:5–37.

